# Variability of quantal NMDA to AMPA current ratio in nucleus tractus solitarii neurons

**DOI:** 10.1101/110569

**Authors:** Caroline Strube, Florian Gackière, Layal Saliba, Fabien Tell, Jean-Pierre Kessler

## Abstract

The ratio between AMPA and NMDA receptors is a key factor governing integrative and plastic properties of excitatory glutamatergic synapses. To determine whether the respective proportions of AMPA and NMDA receptors are similar or vary across a neuron's synapse, we analyzed the variability of NMDA and AMPA currents in quantal responses recorded from neurons located in the nucleus tractus solitarii. We found that the average NMDA to AMPA current ratio strongly differed between recorded neurons and that most of the intra-neuronal current ratio variability was attributable to fluctuations in NMDA current. We next performed computer simulations with a Monte Carlo model of a glutamatergic synapse to estimate the part of AMPA and NMDA currents fluctuations induced by stochastic factors. We found that NMDA current variability mainly resulted from strong channel noise with few influence of release variations. On the contrary, partly because of the presence of subconductance states, AMPA receptor channel noise was low and AMPA current fluctuations tightly reflected changes in the amount of glutamate released. We next showed that these two factors, channel noise and fluctuations in glutamate release, were sufficient to explain the observed variability of the NMDA to AMPA current ratio in quantal events recorded from the same neuron. We therefore concluded that the proportion of AMPA and NMDA receptors was similar, or roughly similar, across synapses onto the same target cell.

## INTRODUCTION

Excitatory glutamatergic synapses in the vertebrate central nervous system (CNS) transmit via two types of ligand gated ion channels, the AMPA and the NMDA receptors. These two types of receptors differ by their pharmacological and biophysical properties. AMPA receptors are low affinity ligandgated channels with fast deactivation whereas NMDA receptors are high affinity receptors with prolonged activation (Traynelis et al., 2010). Consequently, they have different roles. AMPA receptors mainly detect fast glutamate transients whereas NMDA receptors also sense slowly changing and steady state glutamate levels (Yang and Xu-Friedman, 2015). In addition, being highly permeable to calcium ions, NMDA receptors play a key role in activity-induced long term changes in synaptic strength and neuronal excitability. Because of these differences in role and behavior between the two receptor types, the NMDA to AMPA receptor ratio is a key parameter that strongly influences the integrative properties of excitatory synapses. Expression levels of AMPA and NMDA receptor subunits in post synaptic membranes are highly variable and depend on the region investigated, the target neuron type and/or the origin of the fibers that give rise to the presynaptic boutons (Nusser et al., 1998; Nyíri et al., 2003; Shinohara et al., 2008; Tarusawa et al., 2009; Dong et al., 2010; Fukazawa and Shigemoto 2012; Rubio et al., 2014). Furthermore, several forms of synaptic plasticity rely on changes in postsynaptic receptor numbers, especially AMPA receptors numbers, indicating that receptor expression levels at synapses may vary with time and state (Turrigiano, 2000). The factors that determine the relative abundance of AMPA and NMDA receptors in a particular synapse remain largely unidentified. Several studies suggest that the ratio between the two receptors is for a large part a pathway-specific property. In CA1 pyramidal cells for instance, responses from perforant path and Schaffer collateral synapses differ by their AMPA to NMDA charge ratio (Otmakhova et al., 2002). Likewise, cortico-striatal and thalamo-striatal pathways elicit responses with different NMDA/AMPA current ratios in striatal neurons (Smeal et al., 2008; Ellender et al., 2013). Thalamic reticular neurons also receives two types of inputs with different NMDA/AMPA current ratios (Deleuze and Huguenard, 2016). However, these data should be interpreted with caution. As discussed in Myme et al (2003), synaptic responses evoked by electrical stimulation of afferent pathways may fail to provide a reliable view of receptor equipment at synapses. Other studies provide a different view. Recordings performed on hippocampal and neocortical neurons show that the amplitudes of AMPA and NMDA receptor currents are correlated across quantal events recorded from the same cell, suggesting that different synapses onto the same target neuron have a relatively constant ratio of each receptor type (Gompert et al., 1998; Umemiya et al.,. 1999; Watt et al., 2000; Myme et al., 2003; Watt et al., 2004).

The aim of the present study was to determine whether AMPA to NMDA receptor ratio is similar or varies across synapses onto the same neuron. We investigated this question by analyzing the sources of current fluctuations across quantal synaptic responses recorded from a single neuron. Our main objective was to determine whether current ratio variability was high, suggesting heterogeneity of synapses as regards receptor ratio, or low enough to be fully explainable by stochastic factors known to induce current fluctuations at a single synapse (channel noise, variations in vesicular transmitter content). Recording of miniature excitatory post-synaptic currents (mEPSCs) were obtained from retrogradely-identified output neurons of the nucleus tractus solitarii (NTS), a brainstem sensory relay nucleus which receives glutamatergic inputs from visceral afferent fibers via the glossopharyngeal and the vagus nerves and in turn projects onto various brain regions (see Baude et al., 2009 for review). The contribution of stochastic factors to AMPA and NMDA current variability was estimated both by a theoretical approach based on the binomial law and by computer simulations performed using a stochastic synapse model.

## METHODS

Experiments were performed on young (3-6 weeks old) male Wistar rats. All procedures were in agreement with the European Communities Council directive (86/609/EEC).

### Electrophysiological recordings

Recordings were obtained from NTS projections neurons identified by retrograde tracing (Strube et al., 2015). Briefly, young adult rats were anesthetized by an intraperitoneal injection of a mixture of ketamine (50 mg/kg, Imalgène 1000, Centravet, Lapalisse, France) and xylazine (15 mg/kg, Rompun 2%, Centravet) and placed in a stereotaxic apparatus with the incisor bar 2 mm below horizontal. Tracing was performed using either red RetroBeads (undiluted Rhodamine-labeled latex microspheres, Lumafluor Inc., Naples, FL, USA) or Fluorogold (2% in 0.2% saline, Fluorochrome LLC., Denver, CO, USA). Tracer (100 nl) was pressure-delivered through a Hamilton syringe connected to a stainless needle (ID: 0.15 mm, OD: 0.25 mm) at a rate of 1 nl s-1 in the parabrachial nucleus (PBN) or the caudal ventrolateral medulla (CVLM). After wound closure and recovery from anaesthesia, the animals were housed individually. Preparation of medullary slices was made as described before (Balland et al., 2006, 2008; Strube et al., 2015) four to seven days after retrograde tracer injection. For recordings, slices were perfused in a chamber at around 3 ml/min with oxygenated ACSF containing (in mM) 120 NaCl, 3 KCl, 26 NaHCO_3_, 1.25 KH_2_PO_4_, 0.5 ascorbate, 2 pyruvate, 3 myoinositol, 10 glucose, 2.5 CaCl_2_, 2.5 MgCl_2_, 0.02 D-serine and a mixture of GABAA receptors blockers (in *μ*M: 20 bicuculline, 100 picrotoxin) at 32°C. Labeled neurons were visualized using a upright microscope (BX51WI, Olympus Corp., Tokyo, Japan) equipped for fluorescence detection. Whole-cell patch-clamp of NTS neurons were made with an Axopatch 200B (Axon instruments, Foster city, CA, USA), filtered at 2 kHz and digitized at 20 kHz. Series resistance was monitored throughout the experiment and neurons in which this parameter was > 20 MΩ or not stable were discarded. Patch electrodes (2-4 MΩ) contained in mM: 120 cesium methane sulfonate, 10 NaCl, 1 MgCl_2_, 1 CaCl_2_, 10 EGTA, 2 ATP, 0.3 GTP, 10 Glucose, 10 HEPES (pH 7.4). Recordings were performed at +40 mV in order to remove NMDA receptor magnesium block. To record mEPSCs, 1*μ*M TTX was added to the external solution. A computer interfaced to a 12-bit A/D converter (Digidata 1200 using Clampex 9.x; Molecular Devices LLC, Sunnyvale, CA, USA) controlled the voltage clamp protocols and data acquisition.

### Data analysis

Detection of mEPSCs was carried out using the event detection module from the Clampfit software (pClamp, Molecular Device). To prevent any loss of data, detection was performed with two templates, corresponding to events with high or low NMDA/AMPA current ratio respectively, using a loose template match stringency (threshold set to 4). False positives were removed by visual examination of each putative event. A minimum of 30 mEPSCs were collected per neuron. AMPA current amplitude (I_AMPA_) was measured at the peak of the mEPSC (current averaged over 0.3 ms). NMDA current amplitude (I_NMDA_) was obtained by averaging current within a time window starting 5 ms after onset.

We used 5ms duration time windows (as in Watt et al 2000, Hanse and Gustafsson 2001, Myme et al. 2003) since long averaging periods (50 ms) resulted in very high dispersion of data. Measures of variability for I_AMPA_, I_NMDA_ and I_NMDA_/I_NMDA_ ratio were obtained by calculating variances (*σ*) and/or coefficients of variation (CVs). CVs were used when dimensionless comparison was required. Statistical analysis were performed using the Graphpad Instat software.

### Computer simulation

Simulation was performed using a Monte-Carlo model of a glutamatergic synapse (Kessler, 2013). The radii of the axon-dendrite apposition and of the active zone-PSD interface were 500 nm and 200 nm, respectively. No glial membrane or glutamate transporter was included in the model. Glutamate was released in front of PSD center. Depending on the experiment, the number of glutamate molecules released at each synaptic event was either set to 3000 or made variable around a 3000 average value using a Gaussian random number generator. Quantum size was limited by low and high cut-offs set at 1000 and 9000 molecules respectively. Glutamate diffusion was calculated using the equation for Brownian displacement in a three dimensional space:

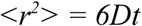

The elementary time step t was set to 10 ns and the coefficient of diffusion for glutamate D was set to 0.4 *μ*m^2^.ms^−1^. AMPA and NMDA receptors were randomly placed in the PSD. NMDA receptors were a mix of GluN2A- and GluN2B-containing receptors (2:8 ratio) to comply with the known presence of GluN2B subunits in NTS NMDA receptors (Zhao et al., 2015). AMPA receptor activation was calculated using the kinetic scheme and rate constants for GluA2-containing receptors from Robert et al. (2005). NMDA receptor activation was calculated using the kinetic scheme 4 from Erreger et al. (2005) and temperature-adjusted rate constants from Santucci and Raghavachari (2008). Temperature correction of NMDA receptor rates was necessary to get rise and decay phases matching those obtained in recording experiments in order to perform measurements in similar conditions. Binding probabilities (*P_on_*) were calculated from association rate constants (*k_on_*) using the following formula:

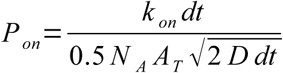

where *N_A_* is the Avogadro number, *A_T_* is the receptor surface area set to 100 nm^2^ and *D* is the diffusion coefficient for glutamate in water (see Kessler 3013, for details). The receptor surface area *A_T_* was used to calculate both collisions of glutamate molecules with receptors and binding probabilities. Thus, it exact value had no incidence on the output of the simulation provided that it was set below an upper limit given by the inverse of the receptor density. The accuracy of binding probability calculation was verified by comparing association curves (without dissociation) obtained by Monte-Carlo methods with those obtained by solving ordinary differential equations using a very simple model consisting in a finite disk (500 nm radius, 12 nm height) populated with 1000 binding sites and 8000 homogeneously dispersed glutamate molecules. Unbinding and transition rates were converted to probabilities using the following general formula :

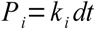

For receptor current calculation, transmembrane potential was set to +40 mV. AMPA receptor conductance was set to 7, 14 and 20 pS for the di-, tri- and quadri-liganded states, respectively. NMDA receptor conductance was set to 50 pS. I_AMPA_ was measured at the peak of the response. Depending on the experiment, I_NMDA_ was either measured 5 ms after glutamate release or obtained by averaging current within a 5 ms duration time window (from 5 to 10 ms after release) in order to match measurements performed on recorded mEPSCs.

## RESULTS

### Variability in mean I_NMDA_/I_AMPA_ ratio across NTS neurons

Recordings were obtained from a total sample of 43 NTS output neurons (see example in Fig. 1A), among which 20 sent projections to PBN and 23 to the CVLM. Data from the two groups of neurons were pooled after checking that there was no significant difference in main mEPSC characteristics according to the projection site (PBN vs CVLM). The frequency of mEPSCs was highly variable ranging from 0.1 to 1 Hz, depending on the neuron (median : 0.22 Hz). At +40 mV, most individual mEPSCs were composite mEPSCs with both a fast and a slow component attributable to AMPA and NMDA receptor activation (I_AMPA_ and I_NMDA_), respectively (Aylwin et al 1997; Balland et al, 2006, 2008, Zhao et al. 2015). We verified that the slow component was suppressed by APV and thus entirely due to NMDA receptor activation (Fig. 1A). Mean I_AMPA_ (*μ*_IAMPA_) exhibited little variability between cells. Depending on the neuron, it ranged from 14 to 27 pA. On the contrary, mean I_NMDA_ (*μ*_I_NMDA__) exhibited five-fold variation across neurons, ranging from 2 to 10 pA. As a consequence, mean I_NMDA_/I_AMPA_ ratio (*μ*_RATIO_) was also highly variable across neurons, ranging from 0.12 to 0.49 (Fig 1B).

**Figure 1.**
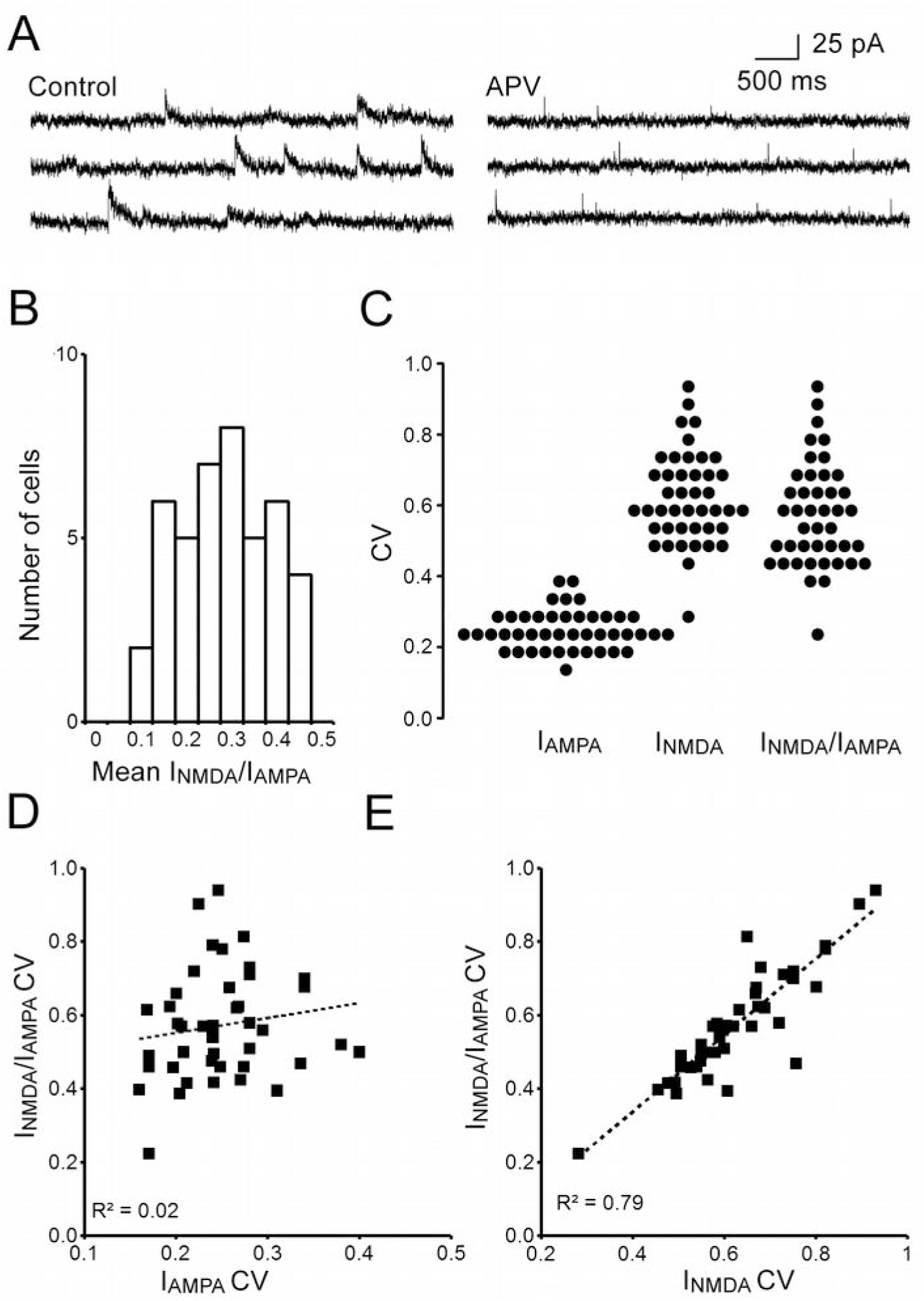
**A.** Example of recordings obtained from an NTS neuron (holding potential + 40mV). The slow component of composites mEPSC recorded in control conditions is no longer present when D,L-APV is added to perfusion medium (100 *μ*M indicating that it is entirely due to NMDA receptors. **B.** Distribution histogram of mean I_NMDA_/I_AMPA_ ratios (μ_RATIO_) in mEPSCs from recorded neurons (n=43). **C**. Distribution histograms of I_AMPA_, I_NMDA_ and I_NMDA_/I_AMPA_ ratio coefficients of variation (CV) in mEPSCs from recorded neurons. **D**. Lack of correlation between I_AMPA_ and I_NMDA_/I_AMPA_ ratio CVs across recorded neurons. **E.** I_NMDA_/I_AMPA_ ratio CV in mEPSCs from recorded neurons linearly increases with I_NMDA_ CV (R^2^ : coefficient of determination).

### Variability in quantal events recorded from the same NTS neuron. Fluctuations of I_NMDA_/I_AMPA_ ratio mainly result from variations of I_NMDA_

To compare the variabilities of I_NMDA_, I_AMPA_ and I_NMDA_/I_AMPA_ ratio across mEPSCs recorded from the same cell we calculated their respective CVs. Intra-neuronal I_NMDA_/I_AMPA_ ratio variability was in some cases relatively high with CV_RATIO_ values up to 0.94 (range 0.22-0.94, depending on the neuron; Fig 1C). We wondered whether this was due to fluctuations in I_AMPA_, or I_NMDA_ or both. Whatever the neuron, CV_IAMPA_ was low ranging from 0.17 to 0.40, indicating little fluctuation from one quantal event to the other (Fig. 1C). I_NMDA_ was far more variable with CV_I_NMDA__ being up to 0.93 and less than 0.4 for 1 neuron only (Fig. 1C). We concluded that fluctuations in I_NMDA_/I_AMPA_ ratio across mEPSCs recorded from the same cell originated from variations in I_NMDA_ rather than variations in I_AMPA_. This finding was confirmed by regression analysis (coefficients of determination : 0.79 versus 0.02, respectively; see Fig. 1D,E). We wondered whether high intra-neuronal variability of I_NMDA_ as compared to I_AMPA_ resulted from differences in receptor channel properties or from stronger variations in NMDA than AMPA receptor content across synapses from the same target cell. To answer this question, we tried to estimate the contribution of stochastic factors to I_NMDA_ and I_AMPA_ variabilities.

### Stochastic factors of I_NMDA_ variability

Two main stochastic factors may contribute to receptor current variability across mEPSCs: random transitions between receptor channel closed and open states (channel noise) and fluctuations in quantal glutamate release. To estimate I_NMDA_ variability resulting from random receptor channel closing and opening, we first calculated channel noise variance according to the binomial distribution. Indeed, if variations of I_NMDA_ across mEPSCs were exclusively due to this factor (i.e. no variation in receptor number, no variation in neurotransmitter quantum size, instantaneous equilibrium of glutamate concentrations within the cleft), then I_NMDA_ would follow a binomial distribution and the resulting variance *σ*^2^_CN_ should be equal to (Sigworth, 1980; Robinson et al., 1991):

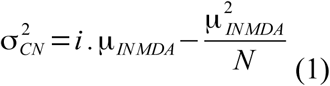

where *N* is the number of NMDA receptors and *i* the unitary receptor current. Since I_NMDA_ is the product of the unitary receptor current *i* by the number of open channels *NP_op_ (P_op_* being the average open probability of NMDA receptors in the synapse), equation 1 may be linearized as follows:

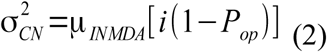

In our experiments, the driving force was set + 40 mV. Thus, *i* was estimated to be about 2 pA (50 pS unitary conductance for GluN2B-containing NMDA receptors, see Traynelis et al., 2010). Assuming a realistic *P_op_* value of 0.1 (Kessler, 2013), we compared I_NMDA_ variances (*σ*^2^I_NMDA_) obtained from recorded neurons with the *σ*^2^_CN_ curve calculated from equation 2 (Fig. 2A). It should be kept in mind that *σ*^2^I_NMDA_ values are likely to have been underestimated since averaging I_NMDA_ measurements over 5 ms duration time-window may have resulted in some smoothing of inter-event fluctuations (see methods). Nevertheless, this comparison suggests that a large part of I_NMDA_ variability across mEPSCS was accounted for by channel noise.

**Figure 2.**
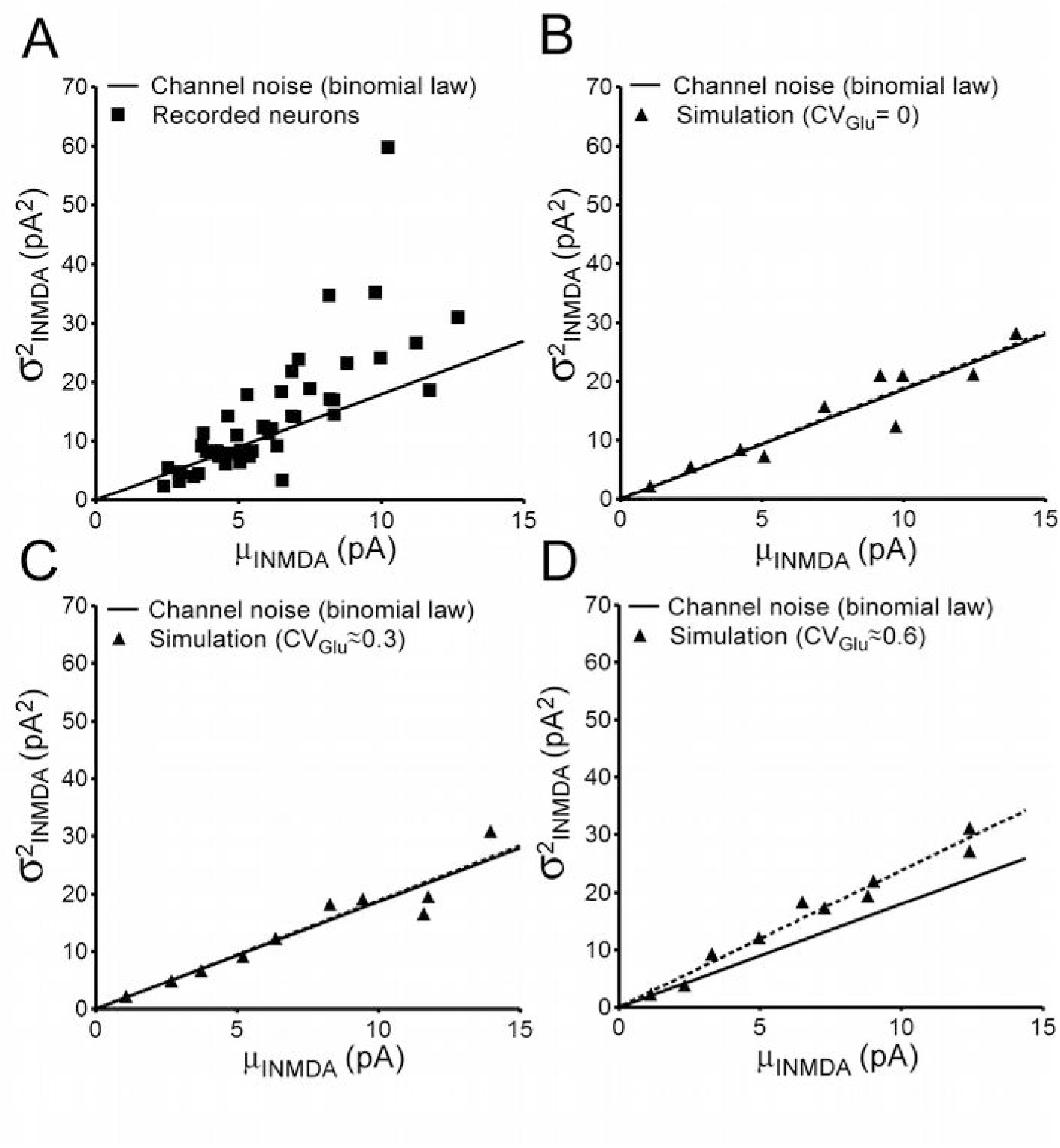
**A. I**_NMDA_ variance (*σ*^2^_INMDA_) across mEPSCs of recorded neurons as a function of mean I_NMDA_. The solid line corresponds to NMDA receptor channel noise values predicted by equation 2 using a 50 pS conductance and a 0.1 P_op_ value. **B**. I_NMDA_ variance in simulated mEPSCs series as a function of mean I_NMDA_. Each data point represents a different mEPSCs series. Data were obtained using a constant amount of glutamate released (3000 molecules per quantal event). Note that the regression line of simulation values (dashed line) perfectly fits with NMDA receptor channel noise values predicted by equation 2 (solid line). **C and D.** As in B, except that the amount of glutamate released (3000 molecules per quantal event on average for each series) was made variable from one quantal event to the other within each simulated mEPSCs series. The coefficient of variation of glutamate released was comprised between 0.26 and 0.31 in C and between 0.50 and 0.62 in D.

Equation 1 relies on the assumption that every NMDA receptor channel in a synapse has the same *P_op_*. This may not be the case since glutamate concentrations decline with distance to the release site. We thus tried to obtain estimates of *σ*^2^_CN_ that take into account possible differences in *P_op_* between receptors according to their location relative to the release site. This was done by computer simulation using a Monte Carlo model of a glutamatergic synapse. Simulation was performed in 10 series of 50 runs each, each run representing a different quantal event. The amount of glutamate released was held constant (3000 molecules) throughout runs and series. The number of NMDA receptors in the synapse was adjusted between series (from 10 to 100) in order to span the entire range of mean I_NMDA_ values obtained from recorded neurons. I_NMDA_ values were measured 5 ms after onset. We compared *σ*^2^I_NMDA_ obtained by simulation with the *σ*^2^_CN_ curve calculated from equation 2 using the average *P_op_* value of NMDA receptors in simulated data (0.07). The fit between the theoretical curve and the simulated data was nearly perfect (Fig. 2B) indicating that binomial distribution based on an averaged *P_op_* provides an accurate description of the stochastic behavior of synaptic NMDA receptors.

We next used computer simulation to get estimate of I_NMDA_ variability resulting from fluctuations in glutamate release. Simulation was performed using a randomly determined amount of glutamate released for each run (see methods). The within-series average was close to 3000 glutamate molecules with either a low (CV_GIU_ ranging from 0.26 to 0.31, depending on the series) or a high (CV_GIU_ ranging from 0.50 to 0.62, depending on the series) variability. Surprisingly, we found little difference between *σ*^2^I_NMDA_ values obtained using either a constant or a randomly varying amount of glutamate release (Fig. 2C,D) suggesting that the part of I_NMDA_ variability resulting from release fluctuations is small as compared to that resulting from channel noise. This finding may seem at odds with the current view which states that channel noise minimally contribute to quantal current variability. This view was mainly based on studies dealing with I_AMPA_ variability (see for instance Franks et al., 2002; 2003). We therefore compared the stochastic behavior of AMPA and NMDA receptors placed in identical conditions.

### Comparison between I_NMDA_ and I_AMPA_ stochastic behavior

Simulation was performed with 100 NMDA receptors and 100 AMPA receptors in the PSD. A first series was obtained with a constant amount of glutamate release throughout runs. Subsequent series were obtained with randomly determined numbers of glutamate molecules released (series average ≈ 3000), using parameters adjusted in order to obtain low, moderate or high release variability (CV_GIU_: 0.3, 0.54 and 0.62, respectively). To allow comparison between I_NMDA_ and I_AMPA_, variances were converted into CVs. We found that contrary to CV_I_NMDA__, CV_IAMPA_ was very low using constant release and steeply increased with CV_GIU_ (Fig. 3A,B). Plotting individual currents values within a series against the amount of glutamate released illustrated the different behaviors of the two receptors (Fig. 3C,D). While I_AMPA_ amplitudes were strongly correlated with release (coefficient of determination : 0.77), I_NMDA_ amplitudes were only loosely correlated with glutamate molecules numbers (coefficient of determination : 0.21). A first factor that may explain this difference is the fact that AMPA receptors had an higher average open probability than NMDA receptors. In addition, AMPA receptors have subconductance states that depend on the number of bound glutamate molecules (Traynelis et al. 2010). It should be kept in mind that equation 2 derives from the binomial distribution and applies to channels that exist in conducting and non-conducting states only, a more complex mathematical description being required for channels with subconductance states, (see Neher and Stevens, 1977). Accordingly, we showed that removing the partially-conducting states (i.e., the di and tri-liganded states) in the AMPA receptor scheme increased I_AMPA_ variability to levels expected from equation 2, indicating that the presence of subconductance states decreases channel noise (Fig. 3E). Noise reduction by subconductance states was substantial as shown by the two-third decrease in variance. Taken as a whole these data point out the fact that, contrary to I_NMDA_ variability which mainly results from receptor noise, I_AMPA_ variability at a single synapse tightly reflects fluctuations in glutamate release. These differences between AMPA and NMDA receptor behaviors may have contributed to the variability of the I_NMDA_ /I_AMPA_ ratio across mEPSCs recorded from the same cell.

**Figure 3.**
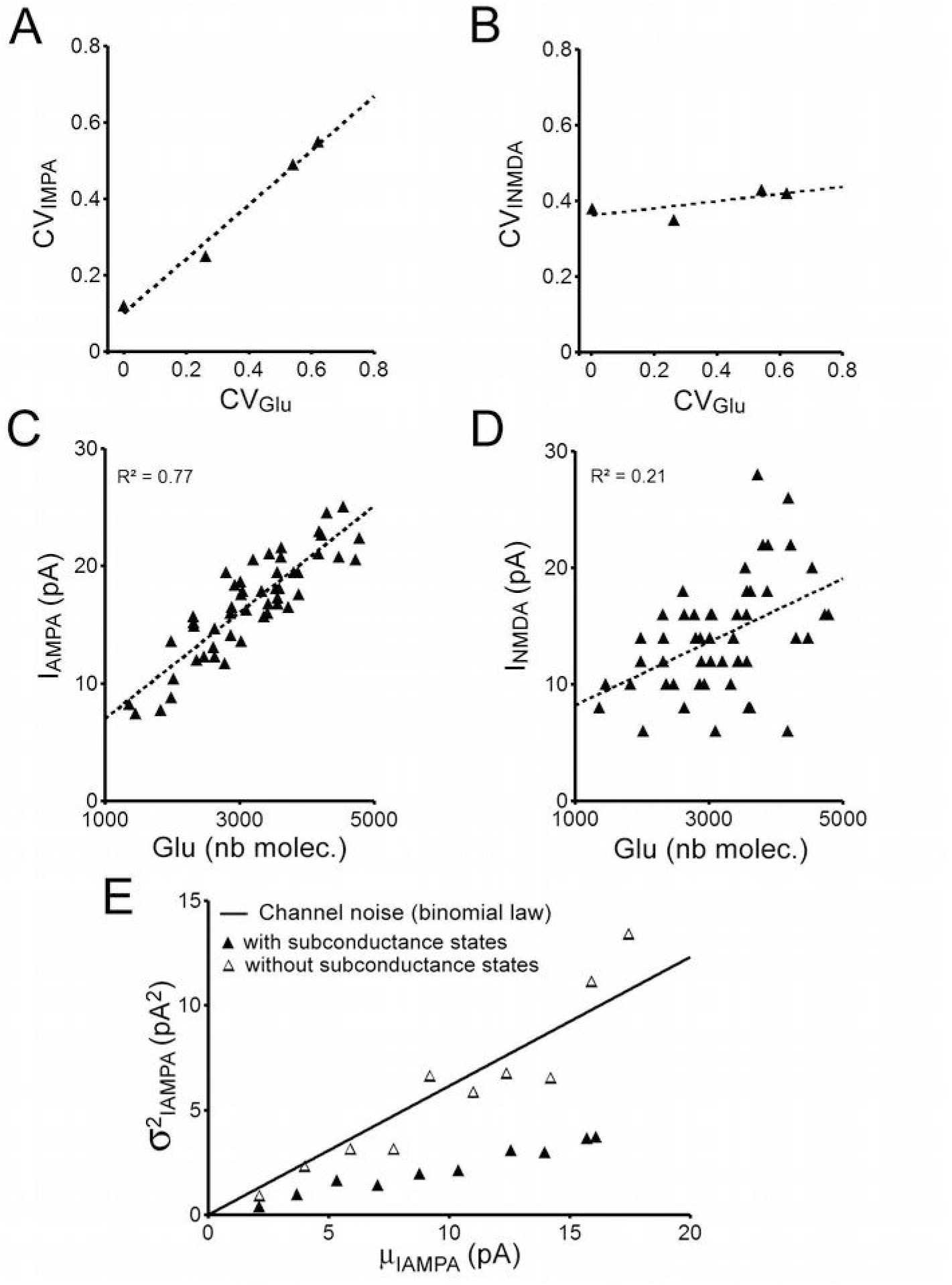
**A and B.** The influence of fluctuations in glutamate release on I_AMPA_ and I_NMDA_ variability. Each data point represents the CV of I_AMPA_ (A) or I_NMDA_ (B) within a series of simulated mEPSCs obtained with 100 AMPA receptors and 100 NMDA receptors and a fixed level of fluctuations in glutamate release. Note the strong correlation between CV_IAMPA_ and CV_GLU_ and the lack of influence of CVGLU on CVI_NMDA_. **C and D.** IAMPA (C) an I_NMDA_ (D) values in individual simulated mEPSCS plotted against the amount of glutamate release. Note that the relationship with the number of glutamate molecules released is strong for I_AMPA_ and considerably weaker for I_NMDA_. **E**. I_AMPA_ variance in simulated mEPSCs series obtained using a constant amount of glutamate released (3000 molecules per quantal event). Each data point represents a different mEPSCs series obtained using either the kinetic scheme from Robert et al. (2005) which includes a 20 pS conductance state and 7 and 14 pS subconductance states (solid triangles) or a simplified kinetic scheme including a single 20 pS conductance state (empty triangles). The solid line corresponds to AMPA receptor channel noise values predicted by equation 2 using a 20 pS conductance and a 0.23 P_op_ value. Note that subconductance states result in decreased I_AMPA_ variance as compared to both expected channel noise and simulation values obtained using the simplified kinetic scheme.

### The origin of of I_NMDA_/I_AMPA_ ratio variability in quantal events recorded from NTS neuron

We next wondered what would be I_NMDA_/I_AMPA_ ratio variability if there were no difference in NMDA/AMPA receptor proportions between synapses onto the same target cell. We estimated *σ^2^_Ratio_* by using first the order Taylor expansion (van Kempen and van Vliet, 2000):

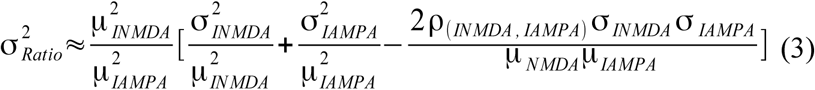

where ρ (I_NMDA_,I_AMPA_) is the correlation coefficient between I_NMDA_ and I_AMPA_. Since I_NMDA_/I_AMPA_ ratio variability was primarily due to variations in I_NMDA_ (see Fig. 1E), we reasoned that receptor ratio heterogeneity across synapses, if present, would primarily result in increased I_NMDA_ fluctuation. Thus, to eliminate potential effects of synapses heterogeneity, we replaced *σ*^2^I_NMDA_ and *σ*I_NMDA_ in equation 3 by *σ*^2^_CN_ and *σ*_CN_ values obtained from equation 2 :

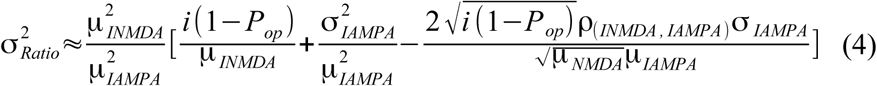

Calculation was performed for each neuron using the experimentally obtained values for *μ*_INMDA_, *μ*_IAMPA_, *σ*^2^_IAMPA_ and *ρ*_(INMDA,IAMPA)_. Comparing *σ*^2^_Ratio_ values directly obtained from recorded data and those recalculated from equation 4 showed that the observed I_NMDA_/I_AMPA_ ratio variability was largely attributable to stochastic factors (Fig. 4A,B). Especially, the slope of the regression line of recorded values on calculated values was close to 1 (Fig. 4B) indicating that experimentally observed variability was on average close to that expected if I_NMDA_ fluctuations were entirely due to channel noise.

**Figure 4.**
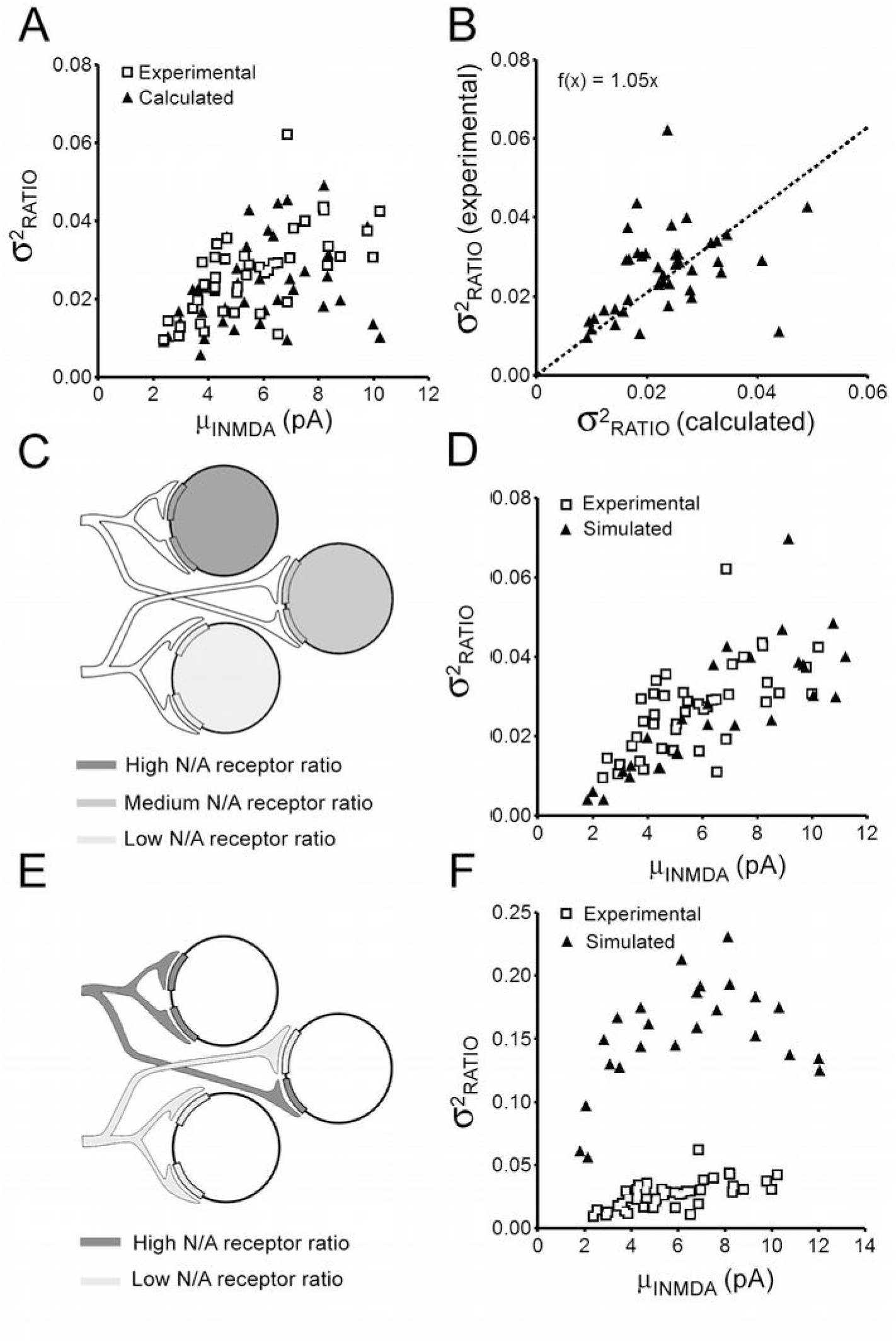
**A.** Variance of I_NMDA_/I_AMPA_ ratio across mEPSCs of recorded neurons as a function of mean I_NMDA_. Note the overlap between experimental data (empty squares) and variances values recalculated for the same neurons using equation 4 (solid triangles). **B.** Regression of experimental ratio variance values on values recalculated using equation 4. The slope of the regression line (origin forced to 0,0) is close to one indicating that I_NMDA_ contribution to ratio variability mas mostly due to NMDA receptor channel noise. **C.** Schematic representation of scenario 1 assuming identical NMDA to AMPA receptor ratio across synapses onto the same target cell. Simulation was performed in 27 series (each representing a different neuron) of 50 runs (each representing a different quantal event). The number of AMPA receptor was set to 100 throughout runs and series. The number of NMDA receptors was identical across runs within a series but increased from 10 to 100 across series. **D.** Comparison between ratio variances obtained from recorded neurons (empty squares) and neurons simulated using scenario 1 (solid triangles). Note the strong overlap between the two sets of data. **E.** Schematic representation of scenario 2 assuming different NMDA to AMPA receptor ratio across synapses onto the same target cell. Simulation was performed in 27 series (each representing a different neuron) of 50 runs (each representing a different quantal event). The number of AMPA receptor was set to 100 throughout runs and series. The number of NMDA receptor was either 5 or 120 depending on the run. The proportion of runs with 120 NMDA receptors increased (from 5:50 to 45:50) across series. **F.** Comparison between ratio variances obtained from recorded neurons (empty squares) and neurons simulated using scenario 2 (solid triangles). Note that ratio variances obtained by simulation using different NMDA to AMPA receptor ratio across synapses onto the same target cell were much higher than those obtained experimentally.

To confirm this finding, we performed simulation according to two scenarios, one assuming that the ratio of NMDA to AMPA receptors at synapses varies between neurons but is strictly identical across the different synapses onto the same neuron (scenario 1, Fig. 4C), the other assuming that the relative abundance of AMPA and NMDA receptors at synapses depends on the afferent pathway only and thus differs between synapses onto the same target cell (scenario 2, Fig 4E). Each neuron was simulated by a series of 50 runs, each run representing a quantal event occurring at a different synapse. The numbers of AMPA and NMDA receptors in the simulation were adjusted in order to fit the averaged quantal currents recorded in actual NTS neurons. We kept the AMPA receptor number constant (100 per synapse) across simulated neurons and we adjusted the overall number of NMDA receptors neuron by neuron in order to span the entire range of *μ*_INMDA_ values found in recorded cells. Release variability was adjusted (CV_GIU_ ≈ 0.3) in order to obtain *σ^2^_IAMPA_* close to those calculated for recorded neurons. I_NMDA_ value was obtained by averaging current over a 5 ms time windows (see methods).

We then plotted *σ^2^_Ratio_* values obtained from either recorded or simulated neurons against *μ*_INMDA_. Variances measured from recorded mEPSCs were very close to values provided by simulation using scenario 1 (Fig. 4D) and far below those obtained using scenario 2 (Fig. 4F), confirming that the variability of the I_NMDA_/I_AMPA_ ratio found in mEPSCs recorded from the same cell resulted from channel noise and fluctuations in glutamate release rather than from heterogeneity of receptor ratio across synapses.

### Correlation between I_NMDA_ and I_AMPA_ across mEPSCs from the same neuron

We wondered whether a similar NMDA to AMPA receptor ratio across a neuron's synapses would invariably result in a strong correlation between I_NMDA_ and I_AMPA_ in mEPSCs. Even with fully-identical receptor ratio, one may expect the correlation to vanish if NMDA channel noise is high enough. Signal to noise ratio (SNR) calculated from the binomial distribution is equal to:

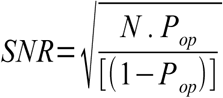

We reasoned that, since SNR increases with the square root of the receptor number *N*, the strength of the correlation between I_NMDA_ and I_AMPA_ should likewise increase with NMDA receptor number. We first look at quantal events simulated using scenario 1. We found a significant correlation between I_NMDA_ and I_AMPA_ for most but not all simulated neurons. Furthermore, the strength of the correlation was highly variable (see examples in Fig. 5A). For the 27 neurons simulated using scenario 1, Pearson r coefficients ranged from 0.14 to 0.65. As expected, correlation strength was found to linearly increase with receptor number in synapses (the only changing parameter between neurons in scenario 1) and hence with *μ*_INMDA_ (Fig. 5B). We next examined recorded neurons. Most but not all (34 out 43) exhibited significant correlation between I_NMDA_ and I_AMPA_ (see example in Fig 5C). Pearson r coefficients ranged from 0.27 to 0.89 and increased with *μ*_INMDA_ (Fig 5D), consistent with the view that loose or lacking correlation resulted from high relative NMDA channel noise rather than synapses heterogeneity.

**Fig 5.**
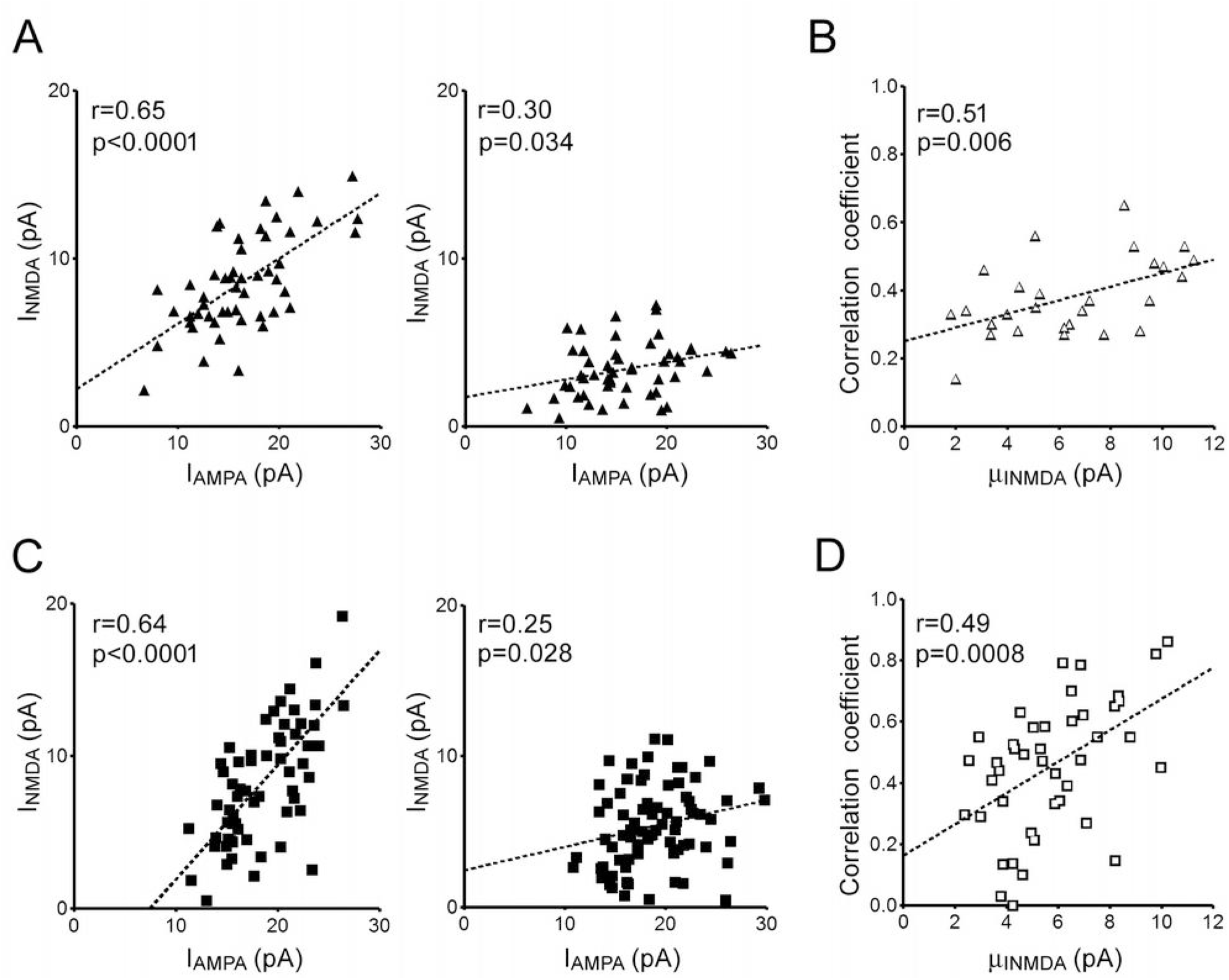
**A.** Example of correlation between NMDA current amplitudes (I_NMDA_) and AMPA current amplitudes (I_AMPA_) across quantal events from two neurons simulated using scenario1. Note the difference in correlation strength between the two neurons. **B.** Relationship between *μ*_INMDA_ and I_NMDA_-I_AMPA_ correlation strength across neurons simulated using scenario 1. **C.** Correlation between NMDA current amplitudes (I_NMDA_) and AMPA current amplitudes (I_AMPA_) across mEPSCs recorded from two NTS projection neurons. **D.** Relationship between *μ*_INMDA_ and I_NMDA_-I_AMPA_ correlation strength across recorded neurons.

## DISCUSSION

Here we found that quantal events recorded from the same NTS projection neuron exhibited substantial variations in I_NMDA_/I_AMPA_ ratio. Using both a theoretical approach and numerical simulation, we showed that variability of I_NMDA_/I_AMPA_ ratio was mostly if not entirely explainable by two factors: i) channel noise being especially strong at NMDA receptors, and ii) fluctuations in glutamate release having stronger effects on I_AMPA_ than on WDA.These findings rule out any substantial contribution of synapse heterogeneity to the variability of I_NMDA_/I_AMPA_ ratio. They therefore imply that the proportions of AMPA and NMDA receptors was similar, or roughly similar, across synapses onto the same target cell. In addition, we found strong differences in mean I_NMDA_/I_AMPA_ ratio between neurons. Thus, our results support the idea that the receptor ratio in synapses is determined by the target cell rather than the afferent pathway. This conclusion is reminiscent of previous findings showing that different synapses onto the same neocortical neuron have very similar NMDA to AMPA receptor ratios (Umemiya et al., 1999; Watt et al., 2000; Myme et al., 2003; Watt et al., 2004) and raises the question of whether mechanisms exist that co-regulate AMPA and NMDA receptor expression in postsynaptic membranes.

As yet, AMPA and NMDA receptor trafficking are viewed as independent processes. To the best of our knowledge, there is no evidence for co-transport of AMPA and NMDA receptors through secretory or endosomal recycling pathways. Likewise, there is no data suggesting that AMPA and NMDA insertion/stabilization in postsynaptic membrane are tightly linked to each other. Alternatively, a conserved receptor ratio across synapses may be the passive consequence of structural constraints. Electron microscope studies performed in various CNS regions using either post-embeding immunogold labeling, freeze fracture replica immunolabeling or STEM tomography indicate that the number of AMPA receptors in synaptic clusters linearly scales with PSD size (Takumi et al., 1999; Racca et al., 2000; Tanaka et al., 2005; Masugi-Tokita et al., 2007; Antal et al., 2008; Shinohara et al., 2008; Dong et al., 2010; Rubio et al., 2014; Chen et al., 2015). NMDA receptor number in synaptic clusters also correlates with PSD size in several brain areas (Racca et al., 2000; Nyiri et al., 2003; Tarusawa et al., 2009; Rubio et al., 2014; but see Takumi et al., 1999; Shinohara et al., 2008; Chen et al., 2015). Thus, it may be hypothesized that the postsynaptic membrane includes finite numbers of specific potential slots for AMPA and NMDA receptors and that the number of slots for each receptor linearly scales with the PSD area. The slot hypothesis was originally proposed to explain how synapses acquire additional AMPA receptors during postsynaptic LTP (Shi et al., 2001; see also Lisman and Raghavachari, 2006; Opazo et al., 2012). It was also postulated that potential slots are not always fully filled with receptors. In this context, a possible interpretation for our data is that the degree of filling of potential NMDA slots is similar across a NTS neuron's synapses but differs between NTS neurons, presumably because of differences in readily available extrasynaptic receptors pools.

An unexpected finding from our simulation experiments was the fact that I_NMDA_ and I_AMPA_ fluctuations across mEPSCs originated from different sources. I_NMDA_ variability was mainly postsynaptic as it resulted from strong channel noise overwhelming the influence of release variations. On the contrary, AMPA receptor channel noise was low and I_AMPA_ variability was mainly presynaptic, tightly reflecting variations in the amount of glutamate released. The lower variability of I_AMPA_ as compared to I_NMDA_ as observed in the present study both *in vivo* and *in silico* is in line with previous results obtained by single synapse recording on hippocampal cell cultures (McAllister and Stevens, 2000). Data in Table 1 from McAllister and Stevens (2000) indicate that the CV of the AMPA component across responses from a single synapse ranged between 0.27 and 0.43, depending on the synapse, while the CV of the NMDA component across the same responses ranged between 0.56 and 0.82, depending on the synapse. It has been claimed previously that differences between the variability of AMPAR and NMDAR responses were due solely to unequal numbers of receptors at the synapse (Franck et al., 2002; 2003). This claim was based on simulations performed with simplified kinetic schemes including few receptor states (Lester and Jahr, 1992 for NMDA receptors and Jonas et al, 1993 for AMPA receptors). Here, using recently published more realistic Markov models (Roberts et al, 2005 for AMPA receptors and Erreger et al. 2005 for NMDA receptors), we unraveled an unexpected biophysical difference between the two receptors. We found that the intrinsic noise of AMPA channels is lower than that of NMDA channels partly as a consequence of AMPA receptors having subconductance states. In addition, the gradual opening of the AMPA receptor pore with the number of bound glutamate molecules provides a mechanism by wich unitary receptor current increases with cleft glutamate concentration. In conclusion, our data show that AMPA receptors are endowed with specific features that reduce the variability of the early as compare to the late NMDA receptor-dependent phase of the postsynaptic response. From a functional point of view, these AMPA receptor specific features may fulfill an important role by increasing the temporal precision and the reliability of fast excitatory transmission.

## Acknowledgments

We wish to think Dr Lydia Kerkerian and Dr Francis Castets for their helpful comments on the manuscript. We also express our gratitude to Dr Boris Barbour for judicious advice in early steps of the study.

## References

Antal M, Fukazawa Y, Eördögh M, Muszil D, Molnár E, Itakura M, Takahashi M, Shigemoto R (2008) Numbers, densities, and colocalization of AMPA- and NMDA-type glutamate receptors at individual synapses in the superficial spinal dorsal horn of rats. J Neurosci 28:9692–96701.

Aylwin ML, Horowitz JM, Bonham AC (1997) NMDA receptors contribute to primary visceral afferent transmission in the nucleus of the solitary tract. J Neurophysiol May;77(5):2539–2548.

Balland B, Lachamp P, Strube C, Kessler JP, Tell F (2006) Glutamatergic synapses in the rat nucleus tractus solitarii develop by direct insertion of calcium-impermeable AMPA receptors and without activation of NMDA receptors. J Physiol. 574:245–261.

Balland B, Lachamp P, Kessler JP, Tell F (2008) Silent synapses in developing rat nucleus tractus solitarii have AMPA receptors. J Neurosci 28:4624–4634.

Baude A, Strube C, Tell F, Kessler JP (2009) Glutamatergic neurotransmission in the nucleus tractus solitarii: structural and functional characteristics. J Chem Neuroanat 38:145–153.

Chen X, Levy JM, Hou A, Winters C, Azzam R, Sousa AA, Leapman RD, Nicoll RA, Reese TS (2015) PSD-95 family MAGUKs are essential for anchoring AMPA and NMDA receptor complexes at the postsynaptic density. Proc Natl Acad Sci U S A 112:E6983–6992.

Deleuze C, Huguenard JR (2016) Two classes of excitatory synaptic responses in rat thalamic reticular neurons. J Neurophysiol. 116:995–1011.

Dong YL, Fukazawa Y, Wang W, Kamasawa N, Shigemoto R (2010) Differential postsynaptic compartments in the laterocapsular division of the central nucleus of amygdala for afferents from the parabrachial nucleus and the basolateral nucleus in the rat. J Comp Neurol 518:4771–91.

Ellender TJ, Harwood J, Kosillo P, Capogna M, Bolam JP (2013) Heterogeneous properties of central lateral and parafascicular thalamic synapses in the striatum. J Physiol 591:257–72.

Erreger K, Dravid SM, Banke TG, Wyllie DJ, Traynelis SF (2005) Subunit-specific gating controls rat NR1/NR2A and NR1/NR2B NMDA channel kinetics and synaptic signalling profiles. J Physiol 563:345–358.

Franks KM, Bartol TM Jr, Sejnowski TJ (2002) A Monte Carlo model reveals independent signaling at central glutamatergic synapses. Biophys J. 83:2333–2348.

Franks KM, Stevens CF, Sejnowski TJ (2003) Independent sources of quantal variability at single glutamatergic synapses. J Neurosci. 23:3186–3195.

Fukazawa Y, Shigemoto R (2012) Intra-synapse-type and inter-synapse-type relationships between synaptic size and AMPAR expression. Curr Opin Neurobiol 22:446–452.

Gomperts SN, Rao A, Craig AM, Malenka RC, Nicoll RA (1998) Postsynaptically silent synapses in single neuron cultures. Neuron 21:1443–1451.

Hanse E, Gustafsson B (2000) Quantal variability at glutamatergic synapses in area CA1 of the rat neonatal hippocampus. J Physiol. 531:467–480.

Jonas P, Major G, Sakmann B (1993) Quantal components of unitary EPSCs at the mossy fibre synapse on CA3 pyramidal cells of rat hippocampus. J Physiol. 472:615–663.

Kessler JP (2013) Control of cleft glutamate concentration and glutamate spill-out by perisynaptic glia: uptake and diffusion barriers. PLoS One 8:e70791.

Lester RA, Jahr CE (1992) NMDA channel behavior depends on agonist affinity. J Neurosci. 12:635–43.

Lisman J, Raghavachari S (2006) A unified model of the presynaptic and postsynaptic changes during LTP at CA1 synapses. Sci STKE 356:re11.

Masugi-Tokita M, Tarusawa E, Watanabe M, Molnár E, Fujimoto K, Shigemoto R (2007) Number and density of AMPA receptors in individual synapses in the rat cerebellum as revealed by SDS-digested freeze-fracture replica labeling. J Neurosci 27:2135–2144.

McAllister AK, Stevens CF 2000). Nonsaturation of AMPA and NMDA receptors at hippocampal synapses. Proc Natl Acad Sci U S A. 97:6173–6178.

Myme CI, Sugino K, Turrigiano GG, Nelson SB (2003) The NMDA-to-AMPA ratio at synapses onto layer 2/3 pyramidal neurons is conserved across prefrontal and visual cortices. J. Neurophysiol 90:771–779

Neher E, Stevens CF. Conductance fluctuations and ionic pores in membranes (1977) Annu Rev Biophys Bioeng. 6:345–381.

Nusser Z, Lujan R, Laube G, Roberts JD, Molnar E, Somogyi P (1998) Cell type and pathway dependence of synaptic AMPA receptor number and variability in the hippocampus. Neuron 21:545–559.

Nyíri G, Stephenson FA, Freund TF, Somogyi P (2003). Large variability in synaptic N-methyl-D-aspartate receptor density on interneurons and a comparison with pyramidal-cell spines in the rat hippocampus. Neuroscience 119:347–63.

Opazo P, Sainlos M, Choquet D (2012) Regulation of AMPA receptor surface diffusion by PSD-95 slots. Curr Opin Neurobiol 22:453–460.

Otmakhova NA, Otmakhov N, Lisman JE (2002) Pathway-specific properties of AMPA and NMDA-mediated transmission in CA1 hippocampal pyramidal cells. J Neurosci 22:1199–207.

Racca C, Stephenson FA, Streit P, Roberts JD, Somogyi P (2000) NMDA receptor content of synapses in stratum radiatum of the hippocampal CA1 area. J Neurosci 20:2512–2522.

Robert A, Armstrong N, Gouaux JE, Howe JR (2005) AMPA receptor binding cleft mutations that alter affinity, efficacy, and recovery from desensitization. J Neurosci 25:3752–3762.

Robinson HP, Sahara Y, Kawai N (1991) Nonstationary fluctuation analysis and direct resolution of single channel currents at postsynaptic sites. Biophys J. 1991 59:295–304.

Rubio ME, Fukazawa Y, Kamasawa N, Clarkson C, Molnár E, Shigemoto R (2014) Target- and input-dependent organization of AMPA and NMDA receptors in synaptic connections of the cochlear nucleus. J Comp Neurol 522:4023–42.

Santucci DM, Raghavachari S (2008) The effects of NR2 subunit-dependent NMDA receptor kinetics on synaptic transmission and CaMKII activation. PLoS Comput Biol. 4:e1000208.

Shi S, Hayashi Y, Esteban JA, Malinow R (2001) Subunit-specific rules governing AMPA receptor trafficking to synapses in hippocampal pyramidal neurons. Cell 105:331–343.

Shinohara Y, Hirase H, Watanabe M, Itakura M, Takahashi M, Shigemoto R (2008) Left-right asymmetry of the hippocampal synapses with differential subunit allocation of glutamate receptors. Proc Natl Acad Sci U S A. 105:19498–19503.

Sigworth FJ (1980) The variance of sodium current fluctuations at the node of Ranvier. J Physiol. 307:97–129.

Smeal RM, Keefe KA, Wilcox KS (2008) Differences in excitatory transmission between thalamic and cortical afferents to single spiny efferent neurons of rat dorsal striatum. Eur J Neurosci 28:2041–2052.

Strube C, Saliba L, Moubarak E, Penalba V, Martin-Eauclaire MF, Tell F, Clerc N (2015) Kv4 channels underlie A-currents with highly variable inactivation time courses but homogeneous other gating properties in the nucleus tractus solitarii. Pflugers Arch 467:789–803.

Takumi Y, Ramírez-León V, Laake P, Rinvik E, Ottersen OP (1999) Different modes of expression of AMPA and NMDA receptors in hippocampal synapses. Nat Neurosci 2:618–24.

Tanaka J, Matsuzaki M, Tarusawa E, Momiyama A, Molnar E, Kasai H, Shigemoto R (2005) Number and density of AMPA receptors in single synapses in immature cerebellum. J Neurosci 25:799–807.

Tarusawa E, Matsui K, Budisantoso T, Molnár E, Watanabe M, Matsui M, Fukazawa Y, Shigemoto R (2009) Input-specific intrasynaptic arrangements of ionotropic glutamate receptors and their impact on postsynaptic responses. J Neurosci 29:12896–12908.

Traynelis SF, Wollmuth LP, McBain CJ, Menniti FS, Vance KM, Ogden KK, Hansen KB, Yuan H, Myers SJ, Dingledine R (2010) Glutamate receptor ion channels: structure, regulation, and function. Pharmacol Rev. 62:405–496.

Turrigiano GG (2000) AMPA receptors unbound: membrane cycling and synaptic plasticity. Neuron. 26:5–8.

Umemiya M, Senda M, and Murphy TH (1999) Behaviour of NMDA and AMPA receptor-mediated miniature EPSCs at rat cortical neuron synapses identified by calcium imaging. J Physiol 521:113–122.

van Kempen GM, van Vliet LJ (2000) Mean and variance of ratio estimators used in fluorescence ratio imaging. Cytometry. 39:300–305.

Watt AJ, van Rossum MC, MacLeod KM, Nelson SB, Turrigiano GG (2000) Activity coregulates quantal AMPA and NMDA currents at neocortical synapses. Neuron 26:659–670.

Watt AJ, Sjöström PJ, Häusser M, Nelson SB, Turrigiano GG (2004) A proportional but slower NMDA potentiation follows AMPA potentiation in LTP. Nat Neurosci. 7:518–24.

Yang Y, Xu-Friedman MA (2015) Different pools of glutamate receptors mediate sensitivity to ambient glutamate in the cochlear nucleus. J Neurophysiol 113:3634–3645.2015

Zhao H, Peters JH, Zhu M, Page SJ, Ritter RC, Appleyard SM (2015) Frequency-dependent facilitation of synaptic throughput via postsynaptic NMDA receptors in the nucleus of the solitary tract. J Physiol; 593:111–125.

